## Supplementary Material for "Interplay between Mg^2+^ and Ca^2+^ at multiple sites of the ryanodine receptor"

**This PDF file includes:**

Figures S1 to S5
Tables S1 to S3
Legends for Movies S1 to S4

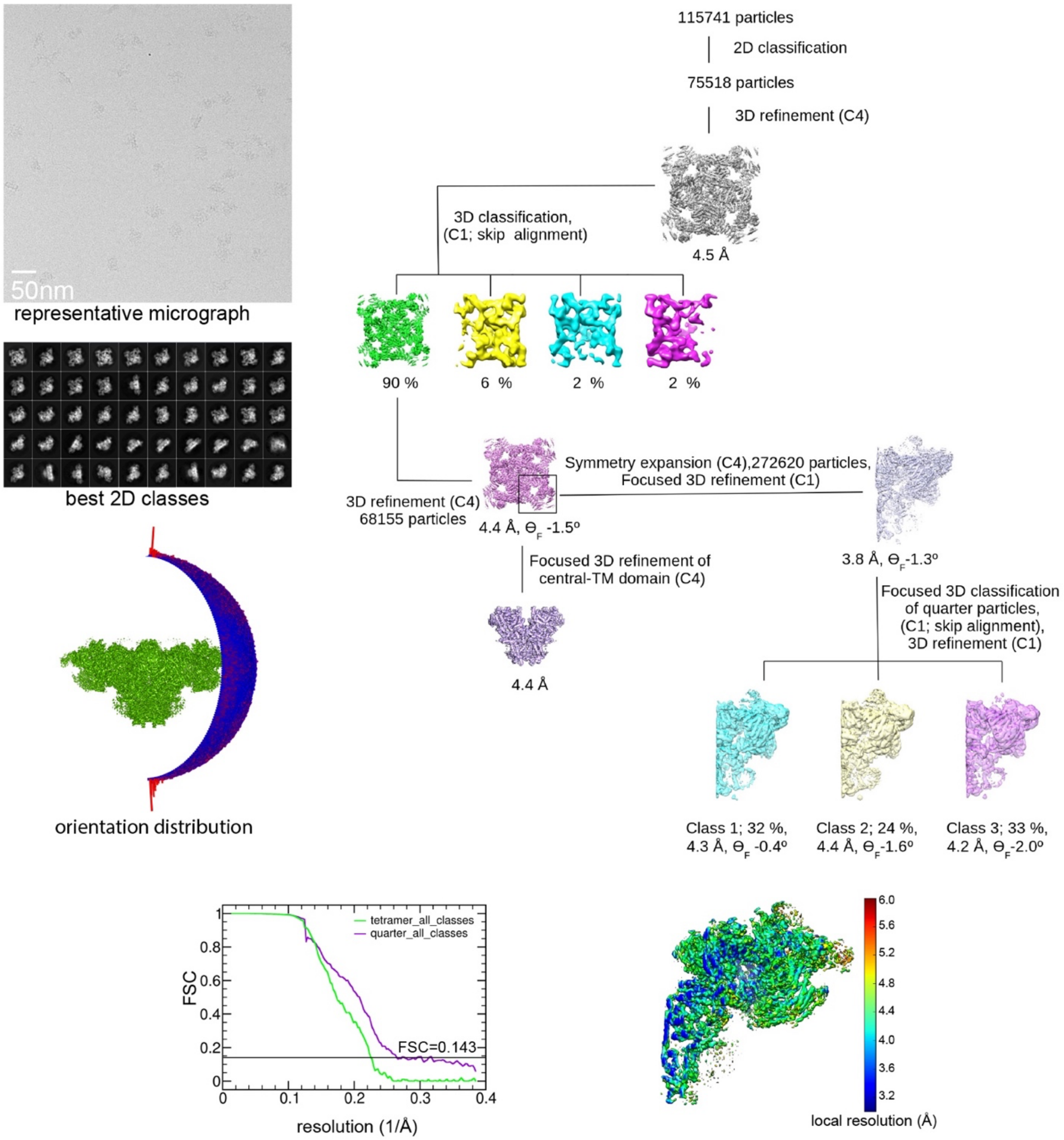


**Figure S1. Image processing scheme for the RyR1-ACP/LM$g^{2+}$ dataset.**

Flowchart depicting a representative micrograph collected on a Titan Krios at 105,000x magnification with a K2 camera in super-resolution mode, 2D class averages obtained by reference-free 2D classification, the entire RyR1 map, and maps focused on the central/transmembrane regions and quarter portion of RyR1. The Euler angle distribution of the particles contributing to the 4.4 Å map, the Gold standard Fourier shell correlation and local resolution of the symmetry-expanded map are also shown. Theta is the flexion angle.

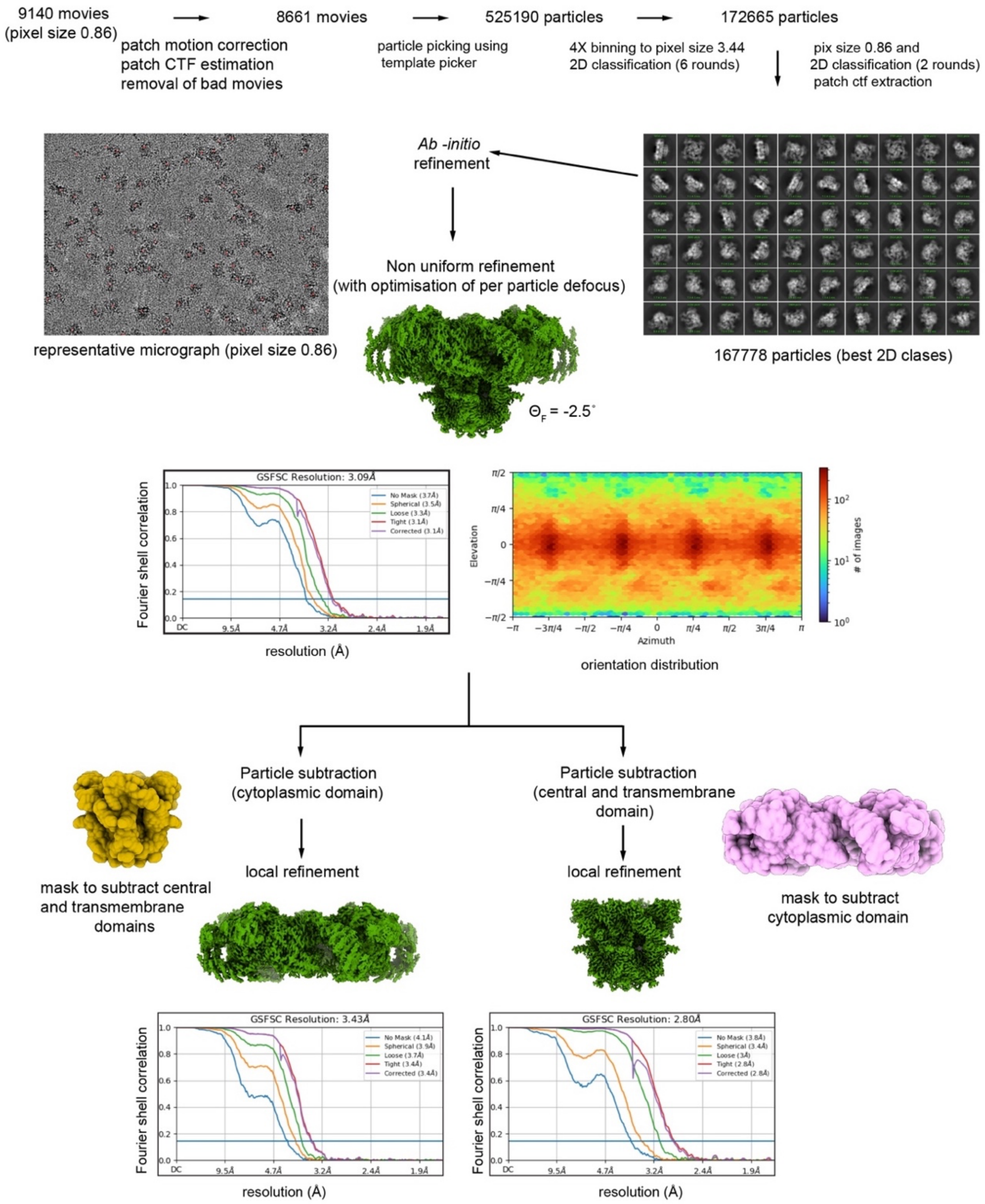


**Figure S2. Image processing scheme for the RyR1-ACP/HMg$^{2+}$ dataset.**

Flowchart depicting a representative micrograph collected on a Titan Krios at 105,000x magnification with a K3 camera in counting mode, 2D class averages after eight rounds of 2D classification, the entire RyR1 map, and maps focused on cytoplasmic and central/transmembrane regions obtained after particle subtraction. The masks used to generate high-resolution reconstructions via particle subtraction and local refinement are shown. The Gold standard (GS) FSC curves of the corresponding cryo-EM reconstructions are depicted. The viewing angle distribution is also shown for the main map. Theta is the flexion angle.

MD replica number 2

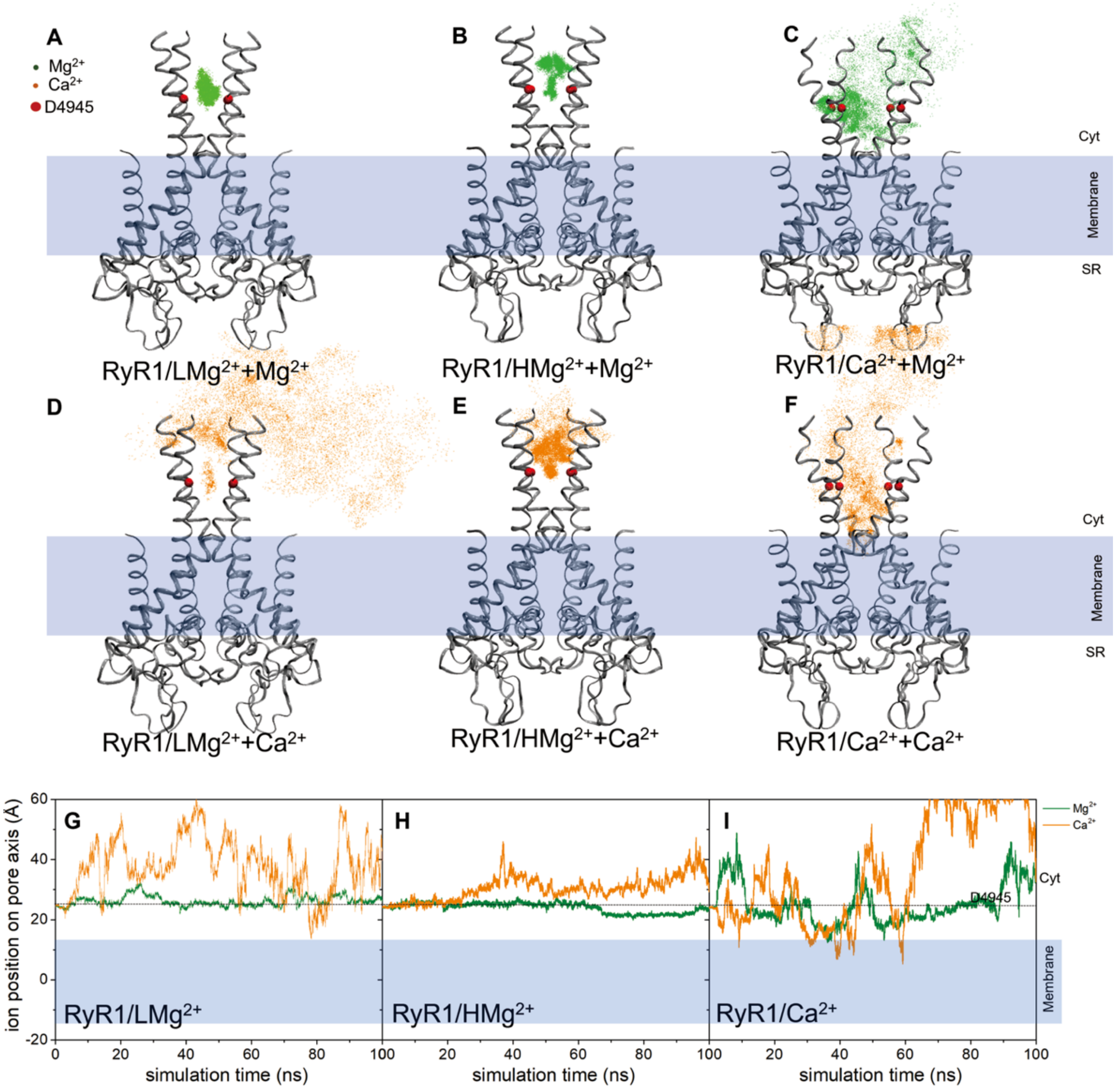

MD replica number 3

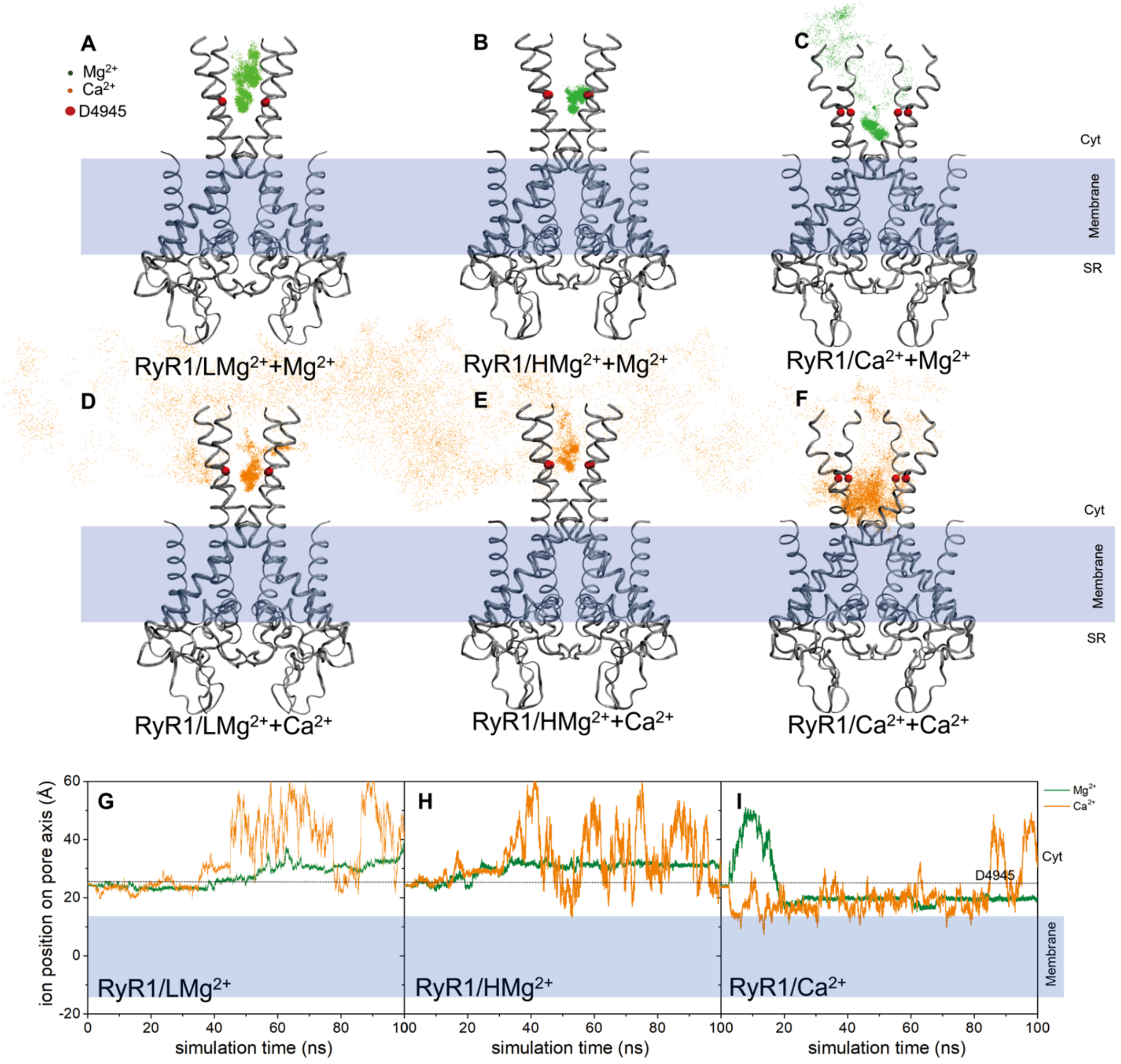


**Figure S3. Replicas of the Molecular dynamics simulations of the pore domain.**

Two replicas illustrating the reproducibility of the MD simulations. **A-F**, Displacement of $Mg^{2+}$ and $Ca^{2+}$ ions is depicted with green and orange dots, respectively, at the D4945 site (red spheres) during 100ns MD simulations. Each dot represents a collection of MD snapshots taken at intervals of 0.02ns. The ribbon structures of the RyR1 pore domain in various conformations are illustrated in their initial configurations. **G-I,** Ion displacement in relation to the z-axis of the channel over the course of simulation time. Note that return to the original z position can occur via the outside of the pore domain. See also Fig. 4.

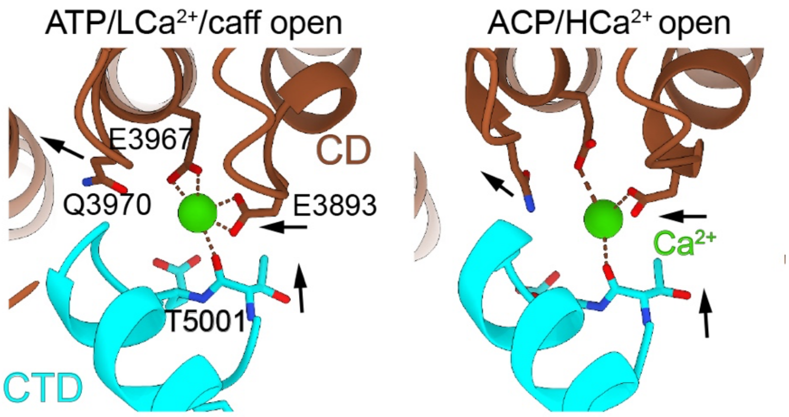


**Figure S4. Binding of $Ca^{2+}$ at the high-affinity $Ca^{2+}$ activation site under open-state conditions.**

The high-affinity $Ca^{2+}$ binding site at the CD/CTD interface has similar configuration at low (30 μM) and high (2 mM) $Ca^{2+}$ concentrations. The PDB IDs are 5TAL and 7TDH, respectively. Arrows indicate the $Ca^{2+}$-induced conformational change. Pore axis is on the left.

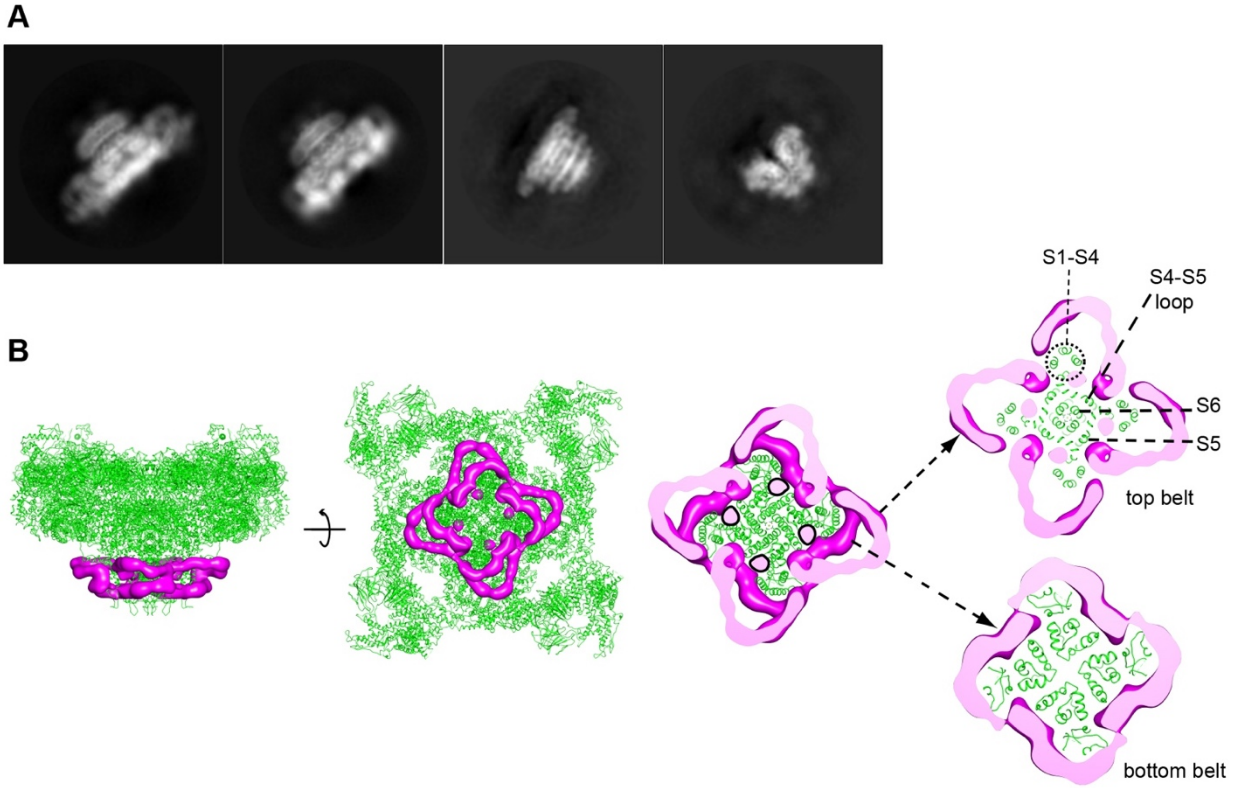


**Figure S5. Nanodisc and lipid densities surrounding the RyR1 TM domain.**

**A**, 2D class averages of the tetrameric map (top left) and CDTM focused map (top right) of RyR1-ACP/LMg$^{2+}$ displaying the nanodisc region surrounding its TM domain. The CDTM-focused reconstruction consists of the CD and transmembrane domains (residues 3668-5037). **B**, Difference map corresponding to the nanodisc obtained by subtracting the model-derived simulated map from the cryo-EM focused map, superimposed to the RyR1-ACP/LMg$^{2+}$ model (left, side view; center, luminal view; right, slices of the TM at different levels of interest). The difference density attributed to the membrane scaffold protein (MSP1E3D1) consists of a top belt that surrounds each voltage sensor-like domain (S1-S4 bundle) plus lipids and a lower smaller belt following more closely the protein density with each segment surrounding one set of voltage sensor-like domains together with S5. Discrete non-protein densities, attributed to lipids, are shown in pink within the nanodisc.

**Table S1. Summary of cryo-EM data collection, image processing and model statistics.**

| | **RyR1-ACP/LMg$^{2+}$** | **RyR1-ACP/HMg$^{2+}$** |
|---|---|---|
| **Data Acquisition** | | |
| Microscope/Detector | Krios/K2 | Krios/ K3 |
| Voltage (kV) | 300 | 300 |
| Magnification | 105000 | 105000 |
| Data collection mode | Super-resolution | Counting |
| Pixel Size (Å) (super-resolution) | 1.32 (0.66) | 0.86 |
| Focus range (μm) | -1 to -2.5 | -1 to -2 |
| Total electron dose (e/Å$^2$) (Number of frames) | 60 (60) | 60 (40) |
| Total number of Micrographs | 3,371 | 9,140 |
| **Image Processing** | | |
| Total number of Particles | 115,741 | 525,190 |
| Particles used (after 3D classification) | 68,155 | 167,778 |
| Resolution (Å) (symmetry expanded) [focused] | 4.4 (3.8) | 3.1 [2.8 TMD, 3.4 CytA] |
| Map sharpening B-factor (Å$^2$) | -250 | - |
| EMDB ID | 22615 | 26610 |
| **Model Refinement** | | |
| RMS Deviation (Bonds) | 0.005 | 0.005 |
| RMS Deviation (Angle) | 0.95 | 1.192 |
| Ramachandran Plot statistics (%) | | |
| Preferred | 91.99 | 93.89 |
| Allowed | 7.78 | 5.62 |
| Outliers | 0.23 | 0.48 |
| **Model Validation** | | |
| Clash-score | 9.66 | 6.34 |
| MolProbity Score | 2.00 | 2.02 |
| PDB ID | 7K0S | 7UMZ |

**Table S2. Simulated systems for MD of the RyR1 Pore Domain[1] in the closed and open states with divalent cations**

| Model ID | PDB ID | State | Cations |
|---|---|---|---|
| **1** | 7K0S | RyR1/LMg$^{2+}$ closed | Mg$^{2+}$ |
| **2** | 7UMZ | RyR1/HMg$^{2+}$ closed | Mg$^{2+}$ |
| **3** | 7TDH | RyR1/Ca$^{2+}$ open | Mg$^{2+}$ |
| **4** | 7K0S | RyR1/LMg$^{2+}$ closed | Ca$^{2+}$ |
| **5** | 7UMZ | RyR1/HMg$^{2+}$ closed | Ca$^{2+}$ |
| **6** | 7TDH | RyR1/Ca$^{2+}$ open | Ca$^{2+}$ |

[1] The Pore Domain encompasses residues 4835-4956; see Methods section for further details.

**Table S3. DFT calculated binding energies of the $[M(H_2O)_n(D4945)_4]^{2+}$ complex of the RyR1 closed structures determined at high and low $Mg^{2+}$ concentrations**

| System | State | $^1E_{AB}$ (Hartree) | $E_A$ (Hartree) | $E_B$ (Hartree) | $\Delta E_{bind}$ (Hartree) | $\Delta E_{bind}$ (kcal/mol) |
|---|---|---|---|---|---|---|
| **Before geometry optimization** | | | | | | |
| $[Mg(H_2O)_6(D4945)_4]^{2+}$ | $HMg^{2+}$ | -2706.099 | -658.527 | -2047.455 | -0.117 | -73.56 |
| $[Mg(H_2O)_6(D4945)_4]^{2+}$ | $LMg^{2+}$ | -2706.068 | -658.517 | -2047.450 | -0.101 | -63.22 |
| $[Ca(H_2O)_7(D4945)_4]^{2+}$ | $HMg^{2+}$ | -3259.941 | -1212.426 | -2047.435 | -0.080 | -50.37 |
| $[Ca(H_2O)_7(D4945)_4]^{2+}$ | $LMg^{2+}$ | -3259.941 | -1212.462 | -2047.419 | -0.061 | -38.13 |
| **After geometry optimization[2]** | | | | | | |
| $[Mg(H_2O)_6(D4945)_4]^{2+}$ | $HMg^{2+}$ | -2706.152 | -658.482 | -2047.519 | -0.152 | -95.10 |
| $[Mg(H_2O)_6(D4945)_4]^{2+}$ | $LMg^{2+}$ | -2706.129 | -658.473 | -2047.509 | -0.147 | -92.13 |
| $[Ca(H_2O)_7(D4945)_4]^{2+}$ | $HMg^{2+}$ | -3259.985 | -1212.366 | -2047.510 | -0.109 | -68.54 |
| $[Ca(H_2O)_7(D4945)_4]^{2+}$ | $LMg^{2+}$ | -3259.985 | -1212.389 | -2047.512 | -0.084 | -52.73 |

Note

[1]$E_{AB}$ : energy of the $[Mg(H_2O)_6(D4945)_4]^{2+}$ or $[Ca(H_2O)_7(D4945)_4]^{2+}$, $E_A$ and $E_B$ : energies of $[Mg(H_2O)_6]^{2+}$ or $[Ca(H_2O)_7]^{2+}$ and $(D4945)_4$, respectively. See Methods section for further details. For the conversion of the energy unit, 1 Hartree ≈ 627.5 kcal/mol.

[2]During DFT optimization, the structure tends to relax to a minimum energy state, which differs slightly from the structure obtained through MD. Notably, the optimized structure of $[Mg(H_2O)_6(D4945)_4]^{2+}$ in $LMg^{2+}$ tends to show a geometry that converges towards that observed in $HMg^{2+}$.

**Movie S1.**

Effect of $Mg^{2+}$ on the pore of RyR1 and tripartite hydrogen bond network.

**Movie S2.**

MD simulation of RyR1/H$Mg^{2+}$+$Mg^{2+}$: stabilization of $Mg^{2+}$ by D4945 depicted in the cytoplasmic view.

**Movie S3.**

MD simulation of RyR1/H$Mg^{2+}$+$Mg^{2+}$: formation of hydrogen bond network among D4945, R4944, D4938, and $Mg^{2+}$, depicted in the side view.

**Movie S4.**

Effect of $Mg^{2+}$ and $Ca^{2+}$ at the high affinity $Ca^{2+}$ activation site and at the EF hand domain, and interrelationship between both domains.
